# Single Chain Fragment Variable (scFv) corresponding to a novel epitope within HCV envelope protein restricts virus entry into hepatocytes

**DOI:** 10.1101/2023.07.07.548124

**Authors:** Soma Das, Padmanava Behera, Dipeshwari J Shewale, Janhavi Bodele, Saumitra Das, Anjali A Karande

## Abstract

Hepatitis C virus (HCV) is a leading cause of chronic viral hepatitis. The use of neutralizing antibodies could be a more effective therapeutic option. Previously we reported the discovery of a novel epitope at the C terminus of HCV-E2 protein, that induced potent neutralizing antibodies in the infected patients. Furthermore, monoclonal antibodies generated against this epitope could also significantly reduce virus replication in a cell culture system. In this study, we have focused on the generation of single chain variable fragments of this unique neutralizing monoclonal antibody A8A11 raised against the conserved epitope. The nucleotide sequence of the neutralizing monoclonal antibody A8A11 was determined and the scFv gene was constructed followed by cloning into the expression plasmid for recombinant protein expression. The scFv mimicked the antibody in binding to the hepatitis C virus like particles (HCV-LP). As expected, the scFv inhibited HCV-LP binding to hepatocytes and could effectively reduce viral replication in the cell culture system. More importantly, scFv A8A11 could restrict serum HCV RNA levels in HCV-infected chimeric mice harboring human hepatocytes. Results provide a basis for developing a promising scFv-based entry inhibitor, which could be more effective against viruses refractory to drugs targeting viral enzymes.

## Introduction

Hepatitis C virus (HCV) represents a global threat to human health. HCV is a blood-borne positive-strand RNA virus belonging to the family Flaviviridae that infects ∼170 million people worldwide (1). 70% - 80% of the infected HCV patients develop chronic hepatitis and 20% - 30% of the chronic patients are found to develop liver disease, cirrhosis and hepatocellular carcinoma (2).

However, therapeutic or protective vaccine is not yet available. Currently used therapies include administration of pegylated alpha interferon in combination with ribavirin (3). The efficacy of HCV treatment has improved by designing multiple direct-acting antivirals (DAAs) based on the recent advances in understanding the lifecycle of HCV. The several classes of DAAs include protease inhibitors, NS5A andNS5 B inhibitors. Several of these inhibitors eg., Teleprevir, Boceprevir, Simeprevir, Ledipasvir, Daclatasvir and Sofosbuvir have been approved recently by the FDA (4). This approach to treatment still suffers a number of drawbacks: regimen restricted to patients with genotype 1, increased rate of adverse effects and low sustained virological response rates (5). Moreover, the effects of the current therapies are likely to be overturned by the emergence of drug resistant viral strains ineffective to replication inhibitors. Therefore, there is a pressing need to develop alternative anti-HCV therapies, particularly in the arena of antibody based therapeutics. The presence of neutralizing antibodies in the serum samples of patients who cleared the infection showed the promise for development of both, prophylactic and therapeutic vaccines. The observation that some HCV-infected individuals (20%) can spontaneously resolve infection with virus-specific immune responses [6] has spurred interest in the potential of HCV vaccines, but as yet no such vaccine exists. Progress toward this goal has been hampered by a number of factors, in particular the extreme genetic diversity of HCV (six major genotypes and more than 50 subtypes) [7]. Identification of protective conserved immune epitopes of the virus is essential for understanding the role of neutralizing responses in disease pathogenesis, and for developing vaccines and antibody-based therapies.

The genetically engineered recombinant antibody fragments have been used in the diagnosis and therapy for treating several diseases. The single chain fragment variable (scFv) is one of the most prevalent types of genetically engineered antibodies [8]. The scFv consists of a variable light chain (V_L_) and heavy chain (V_H_) connected by a short peptide linker. The advantages of scFv are its small size, low immunogenicity, high specificity, and ability to be genetically engineered. The scFv can be produced in bacterial expression systems for large-scale production [9-11]. Several scFvs have been reported to control virus infection, including scFvs against chicken infectious bursal disease virus, scFvs targeting human influenza virus H5N1, and scFvs against the phosphoprotein of Newcastle disease virus [12-14].

Previously using baculovirus-generated HCV-like particles (HCV-LPs) we isolated anti-viral antibodies from patients with acute or chronic HCV infection that were able to specifically inhibit the binding of HCV-LP to hepatic cells. We showed for the first time that neutralizing IgG antibodies generated to the C-terminal region of E2 protein (aa 634–679) successfully reduce HCV genotype 3a infection. More importantly, a stretch of 45 aa (E2C45) encompassing the neutralizing epitopes was designed and mAbs (A8A11) raised against this peptide was able to efficiently inhibit the interactions between HCV LP and hepatocytes as well as *in vitro* HCV infection [15].

A previous study showed the development of an Fv fragment specific for the active site of HCV NS3 protease, but no reports available on the identification of scFvs against HCV envelope protein until date. In this study, we report the nucleotide sequence of the gene segments encoding the variable domains of mAb A8A11, and describe the cloning and expression of a scFv construction corresponding to the antibody [16].

To the best of our knowledge, this is the first report to examine the antiviral activity of scFv against the envelope protein of HCV and our results provide a basis for the development of scFv-based drugs for treatment and prevention of HCV induced disease progression.

## Results and Discussion

To generate the construct for the A8A11 scFvs, total RNA was isolated from A8A11 hybridoma cells which were cultured in Iscove’s modified Dulbecco’s medium (IMDM) supplemented with 10% FBS, antibiotics and 50 μM β-mercaptoethanol. Total RNA was extracted from 10^6^ fresh hybridoma cells using TRIzol reagent (Invitrogen) and used as template to synthesize full-length cDNA using random primers. The variable regions of the heavy and light chains (V_H_ and V_L_) of the A8A11 mAb were amplified by PCR primers (Table 1) and gradient temperature from 45°C, 50°C, 55°C and 60°C (Fig. 1A, as depicted in lanes 1,2,3 and 4). The optimum temperature was determined for HC and LC of A8A11 at 55° C and large-scale PCR was set up to amplify the heavy chain (HC) and light chain (LC) regions of the cDNA. The PCR was found to amplify the expected 340-bp heavy-chain fragment and then the product was cloned into a pGEM-T vector for further sequencing. 4 random colonies were cloned and selected for sequencing and all the colonies showed >99% similarity in sequencing. The full-length V_H_ gene of the A8A11 mAb is 342 bp (Fig. 1C) and can code for a 114-amino-acid protein (Fig. 1D). The same strategy was applied to clone the V_L_ of the antibody (Fig. 1B) and the expected V_L_ gene fragment was purified and cloned into a pGEM-T vector, and then 6 colonies were randomly chosen for sequencing. The data revealed that all of the sequenced clones were identical and contained a 325-bp V_L_ gene fragment (Fig. 1C) that codes for a 112-amino-acid protein (Fig. 1D). The V_H_ gene of A8A11 was 90.28% identical to that of a mouse immunoglobulin heavy chain, NCBI accession no. KU257018), and the V_L_ gene of A8A11was 96.26% identical to that of a mouse immunoglobulin light chain, NCBI accession no. KC422478). The complementarity determining regions (CDRs) within the V_H_ and V_L_ were predicted by use of the IMGT/VQUEST online system (http://www.imgt.org/IMGT_vquest/share/textes/). To generate mAb A8A11 scFv, both the heavy and light chains of variable region of mAb A8A11 were linked together using a polylinker of glycine and serine residues (Gly_4_Ser)3 that is commonly employed for formation of scFvs connecting the C-terminus of V_H_ to the N-terminus of V_L_ (Fig. 1D) and synthesized commercially (Genscript,USA). Finally, the synthesized scFv was cloned in pPROEX HTb vector using EcoRI/HindIII restriction sites. The clone of pPROEX HTb-A8A11 scFv was transformed in Arctic Express cells (Agilent Technologies). Several transformants were inoculated in LB medium (containing 20 μg/ml of gentamycin and 100ug/ml ampicilln) at 37°C and 200 rpm overnight followed by growing the cells without antibiotic selection for 3 hours at 30°C and 200 rpm. Finally, expression of the recombinant protein was induced with 1mM IPTG at 12°C for 24 hours. scFvs frequently form inclusion bodies when they are overexpressed in bacterial cytoplasm. The bacterial cell pellet was resuspended in 20ml of lysis buffer (20mM Tris-HCl pH 8.0, 100mM NaCl, 10mM DTT, 1mM EDTA, 23% (w/v) sucrose, lysozyme (20mg/ml) and DNaseI and incubated for 45min at 37°C on incubator. The suspension was then subjected to ultra-sonication to lyse the cells followed by centrifugation. As evident from Fig. 1E, over expressed protein is present in the inclusion bodies and not secreted in the supernatant, the pellet was subjected to the following steps. Primarily, wash buffer (10mM Tris-HCl pH=8.0, 100mM NaCl, 0.5% Triton X-100, 1mM EDTA, 10mM DTT) was added to the ground pellet (∼15ml) and vortexed vigorously and washed twice. Finally, the pellet was washed with a detergent free wash buffer (10mM Tris-HCl pH=8.0, 100mM NaCl, 1mM EDTA, 10mM DTT). The protein could not be purified in native form, so it was denatured and purified using Ni-NTA. The purified scFv present in different eluted fractions shows a ∼28 kDa band (Fig. 1F) in SDS-PAGE. All the fractions were pooled and refolded by the following protocol. 25mg of protein was diluted to 1mg/ml in guanidine HCl buffer (10mM sodium acetatec, 6M Guanidine hydrochloride, 5mM EDTA, 1mM DTT). Each time 3mg of protein was injected into vigorously stirring refolding buffer (100mM Tris HCl - pH 8.0, 400mM Arginine HCl, 1mM EDTA, 5mM reduced glutathione, 0.5mM oxidized glutathione) and kept on slow constant stirring at 4°C with an interval of 1 hour. The refolded and concentrated protein was confirmed on SDS-PAGE (Fig. 1G) and immunoblot analysis using anti-his antibody further confirmed the presence of scFv (Fig. 1H). To validate the reactivity and binding specificity of scFv against HCV-LP standardized ELISA was utilised. As seen in Fig. 1I, there is significant reactivity between scFv and HCV-LP where 100ng of scFv detected upto 30 ng of HCV-LP. This scFv showed specific binding activity and relatively high affinity for HCV envelope protein, which provided basis for study on inhibition of HCV infection. To detect the inhibitory activity of scFv A8A11 in binding of HCV-LPs to Huh7 cells, different concentration of purified and renatured A8A11 scFv (2.5, 5, 10 and 20 μg/ml) was incubated with Alexa Fluor 488 (Invitrogen) labelled HCV-LPs (gt3a) followed by addition to Huh7 cells (15). The parent monoclonal antibody A8A11 was used as a positive control to compare the inhibitory activity. Binding of HCV-LPs to Huh7 cells was measured by flow cytometry. Cell bound fluorescence was analysed using an FACS calibur flow cytometer (Becton Dickinson) using Cyflogic software to calculate the median fluorescence intensity (MFI) of the cell population, which directly relates to the surface density of Alexa-labelled HCV-LPs bound to hepatocytes. The MFI values of cells with or without HCV-LPs were compared and per cent binding was determined from the equation: % binding of VLP to cells =[(experimental MFI-negative control (only cells) MFI)/(positive control MFI-negative control (only cells) MFI)]X100

**Table 1:**
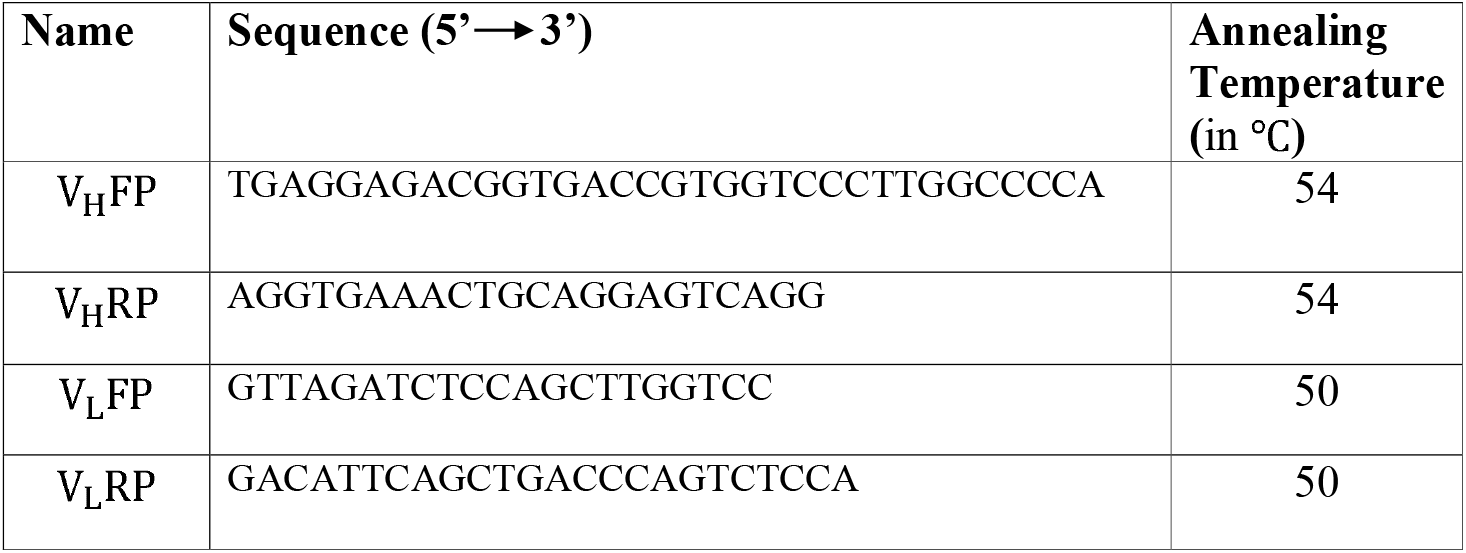
List of Primers used for generation of scFv.

**Figure 1:**
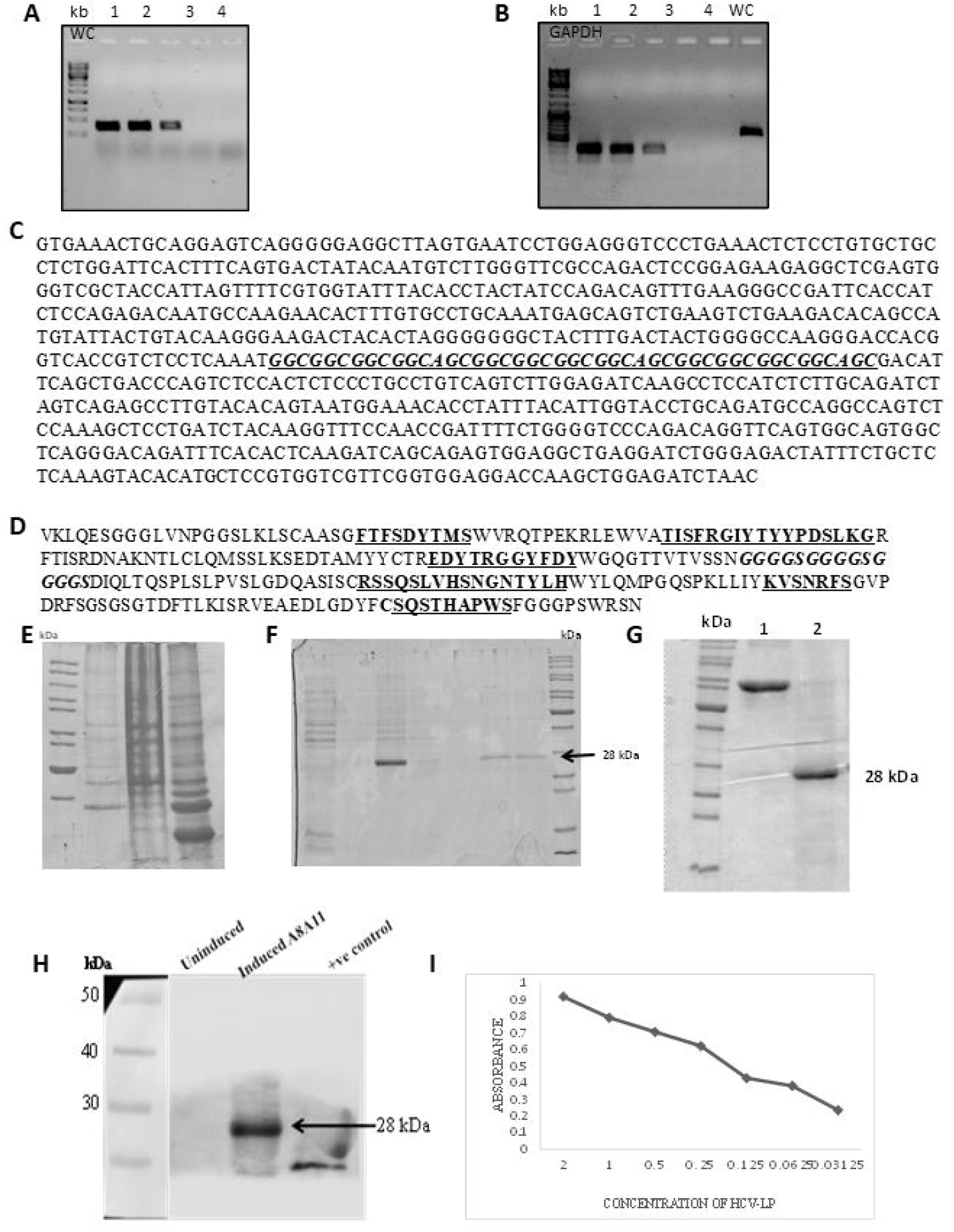
Identification of heavy and light chain of scFv A8A11: PCR amplified product of heavy chain (A) of mAb A8A11 using annealing temperatures as 50°, 55°, 60° and 65° C as seen in lanes 1, 2, 3 and 4 respectively. Similarly, light chain (B) of mAb A8A11 was PCR amplified using the same conditions and annealing temperature. Last lane shows negative control. (C) Nucleotide sequences and (D) amino-acid sequences deduced from the V_H_ and V_L_ regions of the A8A11 hybridoma. CDRs 1, 2, and 3 are indicated in bold and underlined. Amino-acid sequence numbering follows that of Kabat et al. The linker peptide is marked in italics. C-terminus of V_H_ was linked to the N-terminus of V_L_ by peptide linker (G_4_S)_3_. (E) The SDS PAGE shows expression of A8A11 scFv cloned in Arctic Express cells where Lanes M shows the molecular size ladder, Lane 2 shows the lysate of Arctic cells only whereas lanes 3 and 4 depicts the crude lysates of uninduced and induced with 1mM IPTG stained with coomassie brilliant blue R-250. (F) The denatured scFv was purified using Ni-NTA column where lane 1 shows cell lysates of uninduced cells, lane 2 shows the supernatant from induced, lysed cell and lane 3 shows the lysates of induced and lysed pellet. Lane 4 to 7 shows the eluate from NI-Nta columns. (G) The gel picture depicts the refolded scFv protein stained with coomassie brilliant blue R-250. (H) Detection of recombinant A8A11 on SDS-10% PAGE. Lane M indicates protein molecular size standard. Lane 1 is crude lysate of uninduced Arctic cells with the pProEXHtb –scFv clone and Lane 2 is cell lysate induced with IPTG transferred onto nitrocellulose membrane and probed with anti-his antibody developed using DAB substrate. (I) Analysis of antigen binding affinity of A8A11 by ELISA to evaluate the activity of A8A11 to Hepatitis C viral glycoprotein. Binding reactivity of scFv antibody A8A11 with HCV-LP where ELISA was performed with 100ng of scFv and serial double dilutions of HCV-LP. The color was developed with TMB. The data shown are the representative of three similar experiments and are the mean of triplicate samples.

As seen in Fig. 2A, 10 μg/ml concentration of scFv inhibited the binding of the HCV-LPs to Huh7 cells >65 % which is comparable to the full length IgG. Additionally, 20 μg/ml successfully inhibited the cellular HCV-LP binding significantly (∼75 %) to Huh 7 cells. Flow cytometric analysis suggested that the purified scFv A8A11 inhibited HCV-LP binding to Huh7 cells significantly in a dose dependent manner.

**Figure 2:**
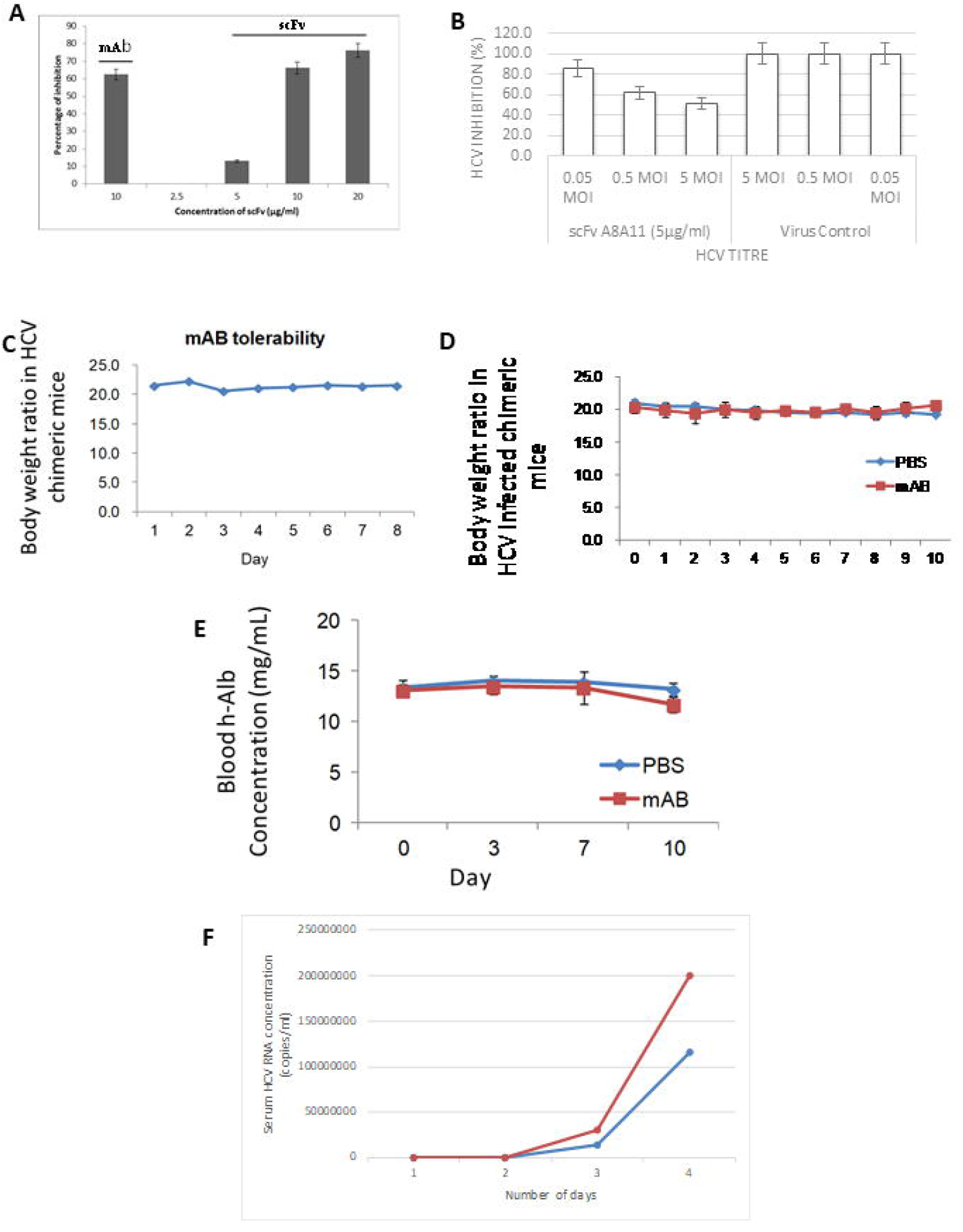
Inhibition of HCV-LP and Huh7 cell binding in HCV infected chimeric mice by purified scFv A8A11: The scFv was pre-incubated with HCV-LPs for 1 h at RT and then the mixture was added to Huh 7cells. HCV-LPs bound to cells was assessed by flow cytometry as described earlier (A) The parent monoclonal antibody A8A11 was used as positive control. The assay was performed in duplicates and carried out at least three times. (B) Inhibition of viral entry in cell culture by the neutralizing scFv A8A11: JFH1 virus of varying titres (0.05 moi, 0.5 moi and 5 moi) was incubated with 5ug/ml of purified scFv A8A11 for 1h at 37°C before infecting Huh 7.5 cells. Three days post infection, total cellular RNA was isolated. HCV negative strand was measured using real time PCR. GAPDH was used as an internal control. The assay was performed in duplicates and carried out at least three times. (C) Tolerability study of scFv A8A11 in chimeric/non PXB mice, n = 2 (with <70% replacement index of primary human hepatocytes). Body weights of these mice were recorded from 0 to 7 days during for repeated administrations of scFv at a dose of 30mg/Kg body weight for 7 days. (D–E) Efficacy study of scFv in HCV infected chimeric/PXB-mice, n = 2 (>70% replacement index of primary human hepatocytes). To study this, uPA/SCID chimeric mice were infected with HCV genotype 3a three weeks prior to challenge with scFv. The body weights and levels of serum h-albumin of post infected PXB mice were monitored daily for 10 days during scFv treatment period (i.e., on 0,3^rd^, 7^th^ and 10^th^ days of infected PXB mice). Similar observations were also recorded in the same time for those post infected PXB mice from control group. (F) Levels of serum HCV RNA titre in control (PBS injected) and scFv treated chimeric/PXB-mice (n = 2).

To verify whether the scFv can also inhibit the binding of virions of hepatitis C, neutralization assays were performed using JFH1 virus as described earlier (17). The construct pJFH1 (a generous gift from Dr Takaji Wakita, National Institute of Infectious Diseases, Tokyo, Japan) was linearized with XbaI followed by transcription of viral RNA. Infectious JFH1 virus particles were generated by transfecting the HCV-JFH1 viral RNA as described previously [17]. Uninfected (mock) Huh7.5 cells were used as control. Huh7.5 cells were seeded in 24-well plates 16 h prior to infection. JFH1 virus was incubated at 37°C for 1 h with serial dilutions of A8A11 scFv. The antibody–virus mixture was then transferred to the cells and infectivity was analyzed 3 days post-infection by qRT-PCR for HCV-negative strand detection [17].

Huh7.5 cells infected with JFH1 virus in the presence of the scFv A8A11 at 5μg/ml showed nearly 85% reduction in intracellular HCV RNA level (Fig. 2B), which was comparable to the inhibition by individual mAb at the same concentration (data not shown).

Finally, to assess the tolerability and *in-vivo* efficacy of the scFv as an effective anti HCV candidate, an inhibition study was carried out in HCV GT 3a-infected PXB mice at Phoenix Bio Inc. Japan.

In order to demonstrate the level of toxicity of scFv, 30mg/Kg body weight of scFv A8A11 was administered by intraperitoneal mode for 7 days to chimeric mice harbouring human liver (engraftment of primary human hepatocytes is around 70–80% in uPA-SCID mice, carried out at Phoenix Bio Inc. Japan). Tolerability result showed that intraperitoneal injection of scFv up to 30mg/Kg body weight for 7 consecutive days is well tolerated in chimeric mice with no signs of toxicity (behavioural changes and mortality) (Figure 2C). Body weights of mice was also found to be normal. After confirming its tolerability in chimeric mice, the therapeutic efficacy of scFv was explored. To study this, first uPA-SCID mice were engrafted with human hepatocytes (> 70% replacement), then these animals were infected with HCV-1b genotype and kept under observation for 3 weeks to reach the HCV RNA titer at optimum levels prior to the challenge study. During this period, the HCV-positive strand RNA copies in the sera of chimeric mice were determined every week by qRT-PCR. Those mice in which serum HCV RNA copy number achieved ≥10^7^, were selected for efficacy study and assigned randomly into two groups of 2 mice each. Group-1 mice were administered with PBS as control. Group-2 mice were treated with scFv protein for 0^th^, 3^rd^ and 7^th^ days. During this window, bodyweight and h-albumin levels, (which is a direct readout of replacement of human hepatocytes into chimeric mice liver), of mice from both groups were recorded. HCV serum RNA from infected mice serum was extracted on 0^th^, 3^rd^, 7^th^ and 10^th^ day of treatment from both the group and quantified by qRT-PCR. The level of HCV RNA copies on 0^th^ day of scFv treatment was approximately ≥10^7^in chimeric mice sera. Subsequently, the treatment of scFv was continued for 3^rd^ and 7^th^ days. The HCV RNA levels in the serum of chimeric mice treated with scFv (group 2) were compared with the chimeric mice treated with PBS (group 1). This experiment was done through contract research with PhoenixBio, Japan.

Significant reduction in HCV viral RNA level was observed in the treated group on 7th and 10th day as compared to the control group, which suggests the potential antiviral activity of scFv A8A11 (Fig. 2F). Results strongly corroborate our previous finding with JFH1 virus (Genotype 2a) in Huh7.5 cells as potent antiviral. Additionally, the levels of human albumin (Fig. 2E) and body weight were measured (Fig. 2D). Results from the studies suggest that HCV chimeric mice were well tolerated upon treatment of scFv at the said concentration, and human albumin levels were also maintained approximately consistent throughout the treatment window. This study provides a proof of concept for the potential of scFv to act as antivirals against HCV.

In summary, the A8A11 scFvs specifically targeting HCV was generated from a neutralizing monoclonal antibody A8A11 which was able to inhibit virus infection in the cell culture system significantly. This scFv showed nontoxicity in vivo and could neutralize hepatitis C virus infection by binding to the E2 protein of HCV. Animal experiments confirmed the protective efficiency of scFv in chimeric mice against HCV challenge. Considering the fact that in spite of DAA therapy, the remaining virus (in some patients) could advance the disease progression to HCC. Use of such scFv based add-on drug might help in eliminating the residual virus and prevent the disease progression more effectively.

## Acknowledgments

This work was supported by the Department of Biotechnology, Ministry of Science and Technology (DBT-IISc Partnership Program); Department of Science and Technology, Ministry of Science and Technology (DST FIST); So. D is supported by WoSA, DST. S.D acknowledges J.C. Bose Fellowship from Department of Science and Technology (DST), Government of India

